# T cell activation and the HLA locus associate with latent infections of human African trypanosomiasis

**DOI:** 10.1101/184762

**Authors:** Paul Capewell, Bruno Bucheton, Caroline Clucas, Hamidou Ilboudo, Anneli Cooper, Taylor-Anne Gorman, Kerry O’Neill, Agapitos Patakas, Andrew Platt, Heli Vaikkinen, William Weir, Mamadou Camara, Paul Garside, Vincent Jamonneau, Annette MacLeod

## Abstract

Infections by many pathogens can result in a wide range of phenotypes, from severe to mild, or even asymptomatic. Understanding the genetic basis of these phenotypes can lead to better tools to treat patients or detect reservoirs. To identify human genetic factors that contribute to symptoms diversity, we examined the range of disease severities caused by the parasite *T. b. gambiense*, the primary cause of human African trypanosomiasis (HAT). We analyzed the transcriptomes of immune cells from both symptomatic HAT cases and individuals with latent infections. Our analysis identified several genes and pathways that associated with the latent phenotype, primarily suggesting increased T and B cell activation in HAT patients relative to latent infections. We also used these transcriptome data to conduct an exome-wide single nucleotide polymorphism (SNP) association study. This suggested that SNPs in the human major histocompatibility locus (HLA) associate with severity, supporting the transcription data and suggesting that T cell activation is a determining factor in outcome. Finally, to establish if T cell activation controls disease severity, we blocked co-stimulatory dependent T cell activation in an animal model for HAT. This showed that reducing T cell activation during trypanosome infection improves symptoms and reduces parasitemia. Our data has used a combination of transcriptome-wide analysis and an *in vivo* model to reveal that T cell activation and the HLA locus associate with the development of symptoms during HAT. This may open new avenues for the development of new therapeutics and prognostics.

## Introduction

Infectious diseases often display a range of severities determined by variation in the pathogen, the host, or both (Riggs et al. 2007; Fakhar et al. 2013; Perron et al. 2008; Lindblade et al. 2014; Hentges et al. 1992). One such medically and economically important disease is human African trypanosomiasis (HAT), primarily caused by the parasite *Trypanosoma brucei gambiense (T. b. gambiense).* Latent infections of *T. b. gambiense* have recently been identified, contradicting more than a century of dogma that the disease is invariably fatal if untreated (Sudarshi et al. 2014; Ilboudo et al. 2011; 2014; Jamonneau et al. 2012). Individuals with latent infections appear able to tolerate the parasite without obvious symptoms for a prolonged period, with one recent case persisting for 29 years without symptoms (Sudarshi et al. 2014). Latent infections are primarily identified using serological methods due to low observable parasitemia. The differences in disease severity found during *T. b. gambiense* infections are likely to be due to host factors, as there is little genetic variation in this clonal parasite (Weir et al. 2016; Kaboré et al. 2011). This provides an opportunity to examine host-derived factors that result in differential disease severity and may provide insight into latent infections with *T. b. gambiense* and possibly other pathogens.

Due to the recent confirmation that latent infections of *T. b. gambiense* exist, there has been limited investigation into potential factors that associate with the phenotype. This work, primarily in a Guinean HAT focus, has demonstrated that elevated IL-8 and IL-6 levels associate with latent *T. b. gambiense* infections. In contrast high levels of immunosuppressive molecules, such as IL-10 and the maternal human leukocyte antigen protein HLA-G, were predictive of disease development in serological positive suspects (Ilboudo et al. 2014; Gineau et al. 2016). Contrasting associations have also been reported with *APOL1* alleles that predispose patients to renal disease (Cooper et al. 2017). The *APOL1* G1 allele was found at higher frequencies in latent infections, while the G2 allele associated with an increased risk of HAT. However, a genome-wide screen has not yet been applied to highlight novel genetic aspects involved in latent *T. b. gambiense* infections. While variations in disease severity caused by African trypanosomes have been extensively studied in mice and cattle, most of these studies used the animal trypanosome species *T. congolense, T. vivax*, and *T. b. brucei* rather than *T. b. gambiense* (Magez et al. 2006; Mansfield and Paulnock 2005; Namangala et al. 2009; Shi et al. 2003; 2006; Tabel et al. 2008). This work has revealed that the immune environment is central to disease outcome in animals. Key features currently elucidated include the involvement of IFNγ and IFNγ-activated macrophages in controlling early infection (Kaushik et al. 1999), the subsequent reduction in the IFNγ-mediated inflammatory environment (Biswas and Mantovani 2014), and the roles that elevated IL-10 and IL-4 play in mediating this shift in the immune environment (Inoue et al. 1999; Mertens et al. 1999).

To more fully understand the immune components involved in disease severity, we compared the peripheral blood mononuclear cell (PBMC) transcriptomes of age-matched groups of *T. b. gambiense* HAT cases, individuals with latent infections, and uninfected controls. Latent infections were defined as individuals who were serologically positive for at least two-years without the development of neurological symptoms or visible parasitaemia using both the card agglutination test for trypanosomiasis (CATT) and the highly *T. b. gambiense* specific trypanolysis test (Jamonneau et al. 2010; Ilboudo et al. 2011). This revealed several differentially expressed genes and pathways associated with disease severity. We also used exome data from the screen to identify single nucleotide polymorphisms (SNPs) that associate with the latent infection phenotype. These data revealed that T cell activation and the major histocompatibility (MHC)/human leukocyte antigen (HLA) locus associate with different disease outcomes in humans. To determine if variation in T cell activation is responsible for variation in disease severity, we blocked co-stimulation dependent T cell activation in a mouse model for trypanosomiasis. This resulted in reduced symptoms (assessed by body weight) and a reduction in parasitemia. These results provided experimental validation of the association with T cell activation identified in the human transcriptomic study, and reveals a central role for T cell activation in determining disease severity in trypanosomiasis.

## Results

### Transcriptome and Pathway Analysis

To identify genetic determinants that underlie disease severity or latent infections, total mRNA was prepared from peripheral blood mononuclear cells (PBMCs) collected from individuals and sequenced using Illumina HiSeq. Transcripts with at least a 1.25-fold difference in expression between HAT patients and latent infection groups and a false discovery rate (FDR)-adjusted p value <0.05, were considered as differentially expressed in this study (Supplemental Table S1). The transcriptional profiles for the HAT and latent infection groups were found to be distinct and could be separated using two-dimensional principal component analysis (PCA) (Figure 1). Additionally, latent individuals clustered more closely with uninfected controls, suggesting that the latent phenotype is due to a lack of response to infection. We found 205 PBMC transcripts (≈0.5%) that significantly differed in abundance (both relatively higher and lower) between HAT cases and latent infections, consisting of 61 transcripts found in greater abundance in individuals with latent infections and 144 that were elevated in HAT cases. There was no enrichment for specific gene ontological (GO) terms, although transcripts greater in abundance in latent infections included regulators of immune response, such as components of the T cell receptor (TCR) complex *(TRAV14, TRAV27)*, IL-4 induced protein 1 *(IL4I1)*, and the gene for the co-stimulatory T cell protein, CD26. *CD8* beta chain transcript was also found to be elevated in latent infections relative to HAT patients. Transcripts greater in abundance in active disease predominantly associate with B cells (primarily immunoglobins and *CD38).* This may reflect the large B cell expansion that has been previously observed in clinical trypanosomiasis (Lejon et al. 2014).

**Figure 1.**
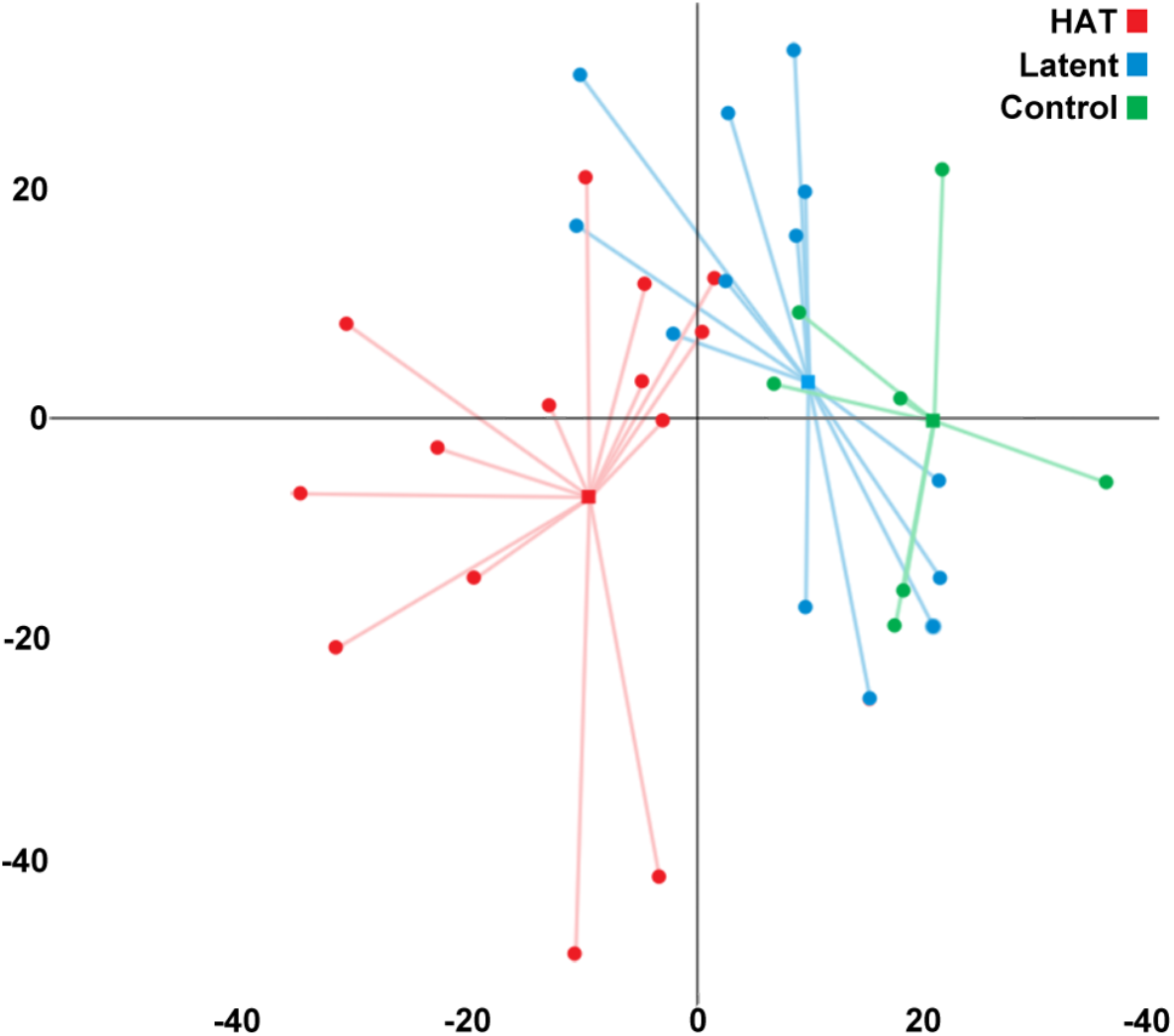
Principle component analysis (PCA). A PCA describing the variability present in the expression profiles of clinical cases, (blue) latent infections (red), and uninfected controls (green). Lines join each sample (circle) to the centroid (square) of the respective group. The two dimensions describe 26% of the variability present in the transcript profiles of the samples.

The biological functions and pathways that may differ due to the different transcriptional profiles found in active disease and latent infections were investigated using Ingenuity Pathway Analysis and a database of experimentally-verified protein functions. This indicated that 102 functional pathways differ between the HAT cases and individuals with latent infections, although the likely direction of regulation between groups (those with a z-score <2.0 or 2.0>) could only be predicted for one pathway, Ca^2^+ mobilization (z-score 2.158, p=0.0007). This would suggest increased lymphocyte activation in HAT patients, relative to latent cases. Other significant pathways (p<0.01) suggest involvement of the binding, activation and proliferation of B and T lymphocytes. Ingenuity Pathway Analysis specifically predicted that activation of lymphocytes (p<0.0001), activation of T cells (p=0.0004), co-stimulation of T lymphocytes (p=0.0013), and Ca^2^+ flux (p=0.0004) (indicative of lymphocyte activation) were all significantly different between HAT patients and latent cases (Supplemental Table S2). Interestingly, pathways associating with an immune response in the skin and joints, including atopic dermatitis (p=0.008), psoriatic arthritis (p=0.006), and inactivation of mast cells (p=0.005), were also implicated. The skin has recently been shown to be an overlooked anatomical reservoir for *T. b. gambiense* (Capewell et al. 2016).

Further Ingenuity Pathway Analysis identified 19 predicted upstream effectors that could explain the expression profile differences between HAT cases and latent infections, including six transcriptional regulators (GATA2, SP1, GATA1, SATB1, CREM, and IZF1)( Supplemental Table S3). Of these, only the SATB1 pathway had a significant z-score that was predicted to be inhibited in latent cases relative to HAT patients (z score=-2.093, p=0.008). The activity of the other five transcription factors could not be predicted from our data. Other upstream putative effectors that may explain the expression differences between HAT and latent infections include several immune regulators, such as CD40 (p=0.001), IL-4 (p=0.002), CXCL12 (p=0.004), and IL-21 (p=0.040). These pathways are each associated with CD4+ T cell activation and predicted to be less active in latent individuals, relative to HAT cases (Supplemental Table S3). CD40, IL-4, and IL-21 are also potent mediators of B cell proliferation and immunoglobulin release (Eto et al. 2011; Spriggs et al. 1992). Supporting a prior study (Gineau et al. 2016), HLA-G (p=0.004) was also an upstream predicted effector for the observed differences in transcriptional profiles, although the predicted direction was unable to be determined by Ingenuity Pathway Analysis. In summary, these data indicate that individuals with latent infections demonstrate a reduced B and T cell response, relative to patients diagnosed with active disease.

### SNP Association

To identify a potential genomic basis for the differences in disease outcome, single nucleotide polymorphisms (SNPs) were identified from the exome data. In total, 39,897 high quality SNPs were identified and their potential associations with HAT and latent infections were examined (figure 2). This highlighted 482 SNPs that differed between individuals with an active or latent infection using a false discovery rate (FDR)-adjusted significance value of p < 0.05 (Supplemental Table S4). fastSTRUCTURE analysis of the marginal likelihoods for various values of K indicated that the most likely number of populations was 1 (marginal likelihood = -0.45), suggesting there was no underlying substructuring that would affect subsequent analysis. The most significant latent infection-associating SNP was found on exon 5 of the class I major histocompatibility complex (MHC) gene, *HLA-A* (adjusted p=0.00089). This polymorphism causes a nonsynonymous change at the junction of the *HLA-A* α3 CD8-binding region and membrane-addressing domains. The majority (71 %) of individuals with latent infections possess this SNP allele compared to 31 % of HAT cases, equating to an estimated odds ratio of 5.231. The statistical power to detect an effect of this size with α=0.05 is 0.6 for a study of this size. Four additional significantly associating SNPs were also found in the HLA-A α3 domain and the neighboring *HLA-B* gene. A second locus of high association was found in a region containing several tripartite motif-containing (TRIM) genes that are involved in retroviral defense (Figure 2)(Nisole et al. 2005). Examining the distribution of SNPs on a gene-by-gene basis revealed no association between individual *TRIM* genes and disease outcome, but there was a significant association with *HLA-A* (adjusted nominal p=0.02597) (Supplemental Table S5).

**Figure 2.**
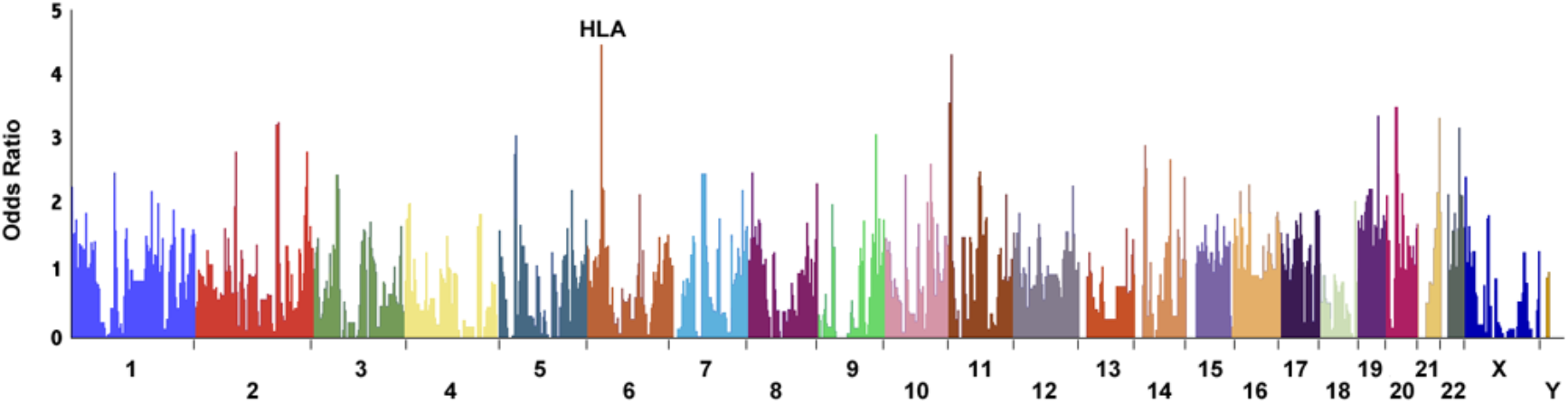
The segmented mean odds ratios of the association between SNPs and the latent infection phenotype. The means odd ratio of genetic association with each SNP was calculated using PLINK. Each presented segment includes a genomic region encompassing two neighboring SNPs, with the mean value determined from the odds ratios of the two SNPs. The highest peak of association lies within the HLA region on chromosome 6.

### The impact of blocking co-stimulation dependent T cell activation on disease outcome in an animal model of trypanosomiasis

Our data from human infections indicate that T cell activation in individuals with latent infections is lower than those experiencing active disease, implying a role for trypanosome-induced T cell activation in disease outcome in trypanosomiasis. In order to test this hypothesis, we used a specific inhibitor of co-stimulation dependent T cell activation (CTLA4-Ig) in the mouse model of symptomatic disease (Linsley et al. 1992). Balb/c mice were either treated with CTLA4-Ig or an isotype control, administered intraperitoneally the day before parasite inoculation and every other day after infection for the length of the experiment. Mice were infected with the trypanosome genome reference strain, TREU927, and the expression of T cell activation markers, parasitemia, and weight loss (an indicator of animal health) were measured and compared to uninfected control mice. CTLA4-Ig significantly reduced the expression of activation-associated immune checkpoint proteins (ICOS, PD1, LAG3) in the CD8+ and CD4+ T cell populations during infection (Supplemental Figure S1). Trypanosome infections typically follow a sinusoidal pattern through disease progression, although we found that reducing T cell activation led to a reduction in parasitemia after the initial peak of parasitaemia (GLM, F statistic 12.96, p<0.001) and an associating recovery in animal condition (assessed using body weight) (ANOVA, p=0.025) (Figure 3). This suggests that co-stimulation dependent T cell activation promotes survival of the parasite in the host and interfering with this biological pathway can induce a less severe disease phenotype in an animal model, mirroring the human latent infection phenotype.

**Figure 3.**
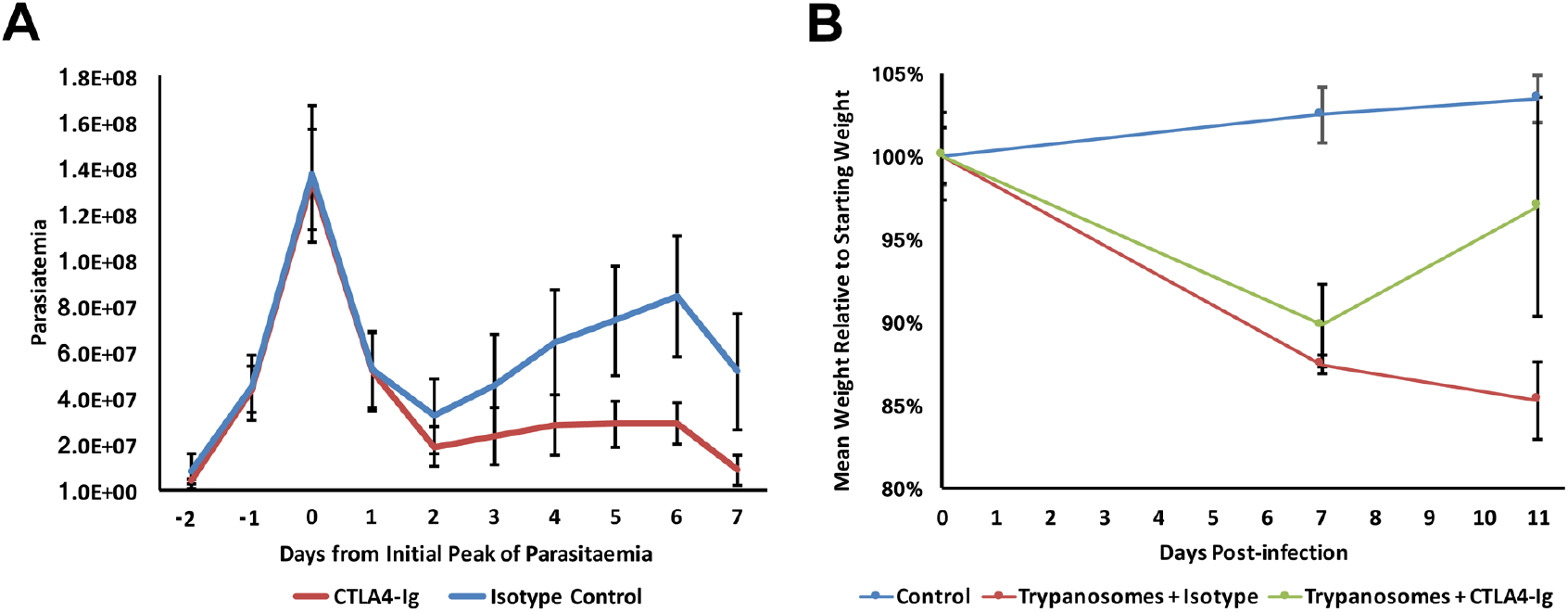
The effects of reducing co-stimulation dependent T cell activation during trypanosome infection. (a) CTLA4-Ig treatment significantly reduces parasitemia after the first peak of parasitemia in trypanosome BALB/c mice infected with *T. brucei* strain TREU 927 (red) versus mice administered an isotype control (blue) (GLM, F statistic 12.96, p<0.001). Parasitemia from different mice are aligned to the first peak to account for variation parasitemia curves (0 = first peak of parasitemia). Data represents fifteen mice in each group with a 95% confidence interval. (b) Mean weight change from day 0 of five mice that were either uninfected (blue), infected with *T. brucei* strain TREU927 and administered an isotype control (green) or infected with *T. brucei* strain TREU927 with regular administrations of CTLA4-Ig (red). In all experiments, CTLA4-Ig was administered intraperitoneally the day before inoculation and every other day after, for the length of the experiment.

## Discussion

Latent infections of *T. b. gambiense* have only recently been identified and there has been little investigation into the genetic factors that may play a role in determining disease outcome in such individuals (Sudarshi et al. 2014; Ilboudo et al. 2011; 2014; Jamonneau et al. 2012). These latent infections likely represent an overlooked and hidden reservoir that will hamper ongoing control efforts for African trypanosomiasis. Interestingly, our study found that pathways involved in a skin immune response significantly differed between active and latent cases. It has recently been demonstrated that extravascular *T. b. gambiense* skin-dwelling parasites can form an overlooked anatomical reservoir in experimentally infected mice and humans living in HAT endemic areas. Individuals harboring latent infections may therefore form an overlooked anatomical reservoir, despite an absence of blood parasites and HAT specific symptoms, such as neurological signs (Capewell et al. 2016). This suggests a putative link between the host immune response and progression to a latent phenotype that requires further examination. To understand the underlying mechanisms that may contribute to the diversity in trypanosomiasis disease severity, our study first examined the transcriptomic differences between latent individuals and clinical cases. There were significant differences between the transcription profiles of latent infected and active cases. Further examination using Ingenuity Pathway Analysis highlighted activation and proliferation of lymphocytes as significantly different pathways, with a suggestion through they were less active in latent infections relative to HAT cases based on Ca^2^+ mobilization. We next used exome-data to identify SNPs that associate with the latent infection phenotype, albeit with low power due to a limited sample size. This suggested an association with the HLA locus, a region directly involved in T cell activation, with the latent and clinical phenotypes. Interestingly, no association was found with genes encoding proteins known to be involved in the human infectivity of African trypanosomes (APOL1, HPR (Vanhollebeke et al. 2008; Vanhamme et al. 2003)). Indeed, *APOL1* alleles have previously been found to associate with the development of a latent phenotype in this population (Cooper et al. 2017). This discrepancy may a result of using a population comprised of immune cells that do not express these proteins or simply due to population size.

The SNPs putatively identified in our study may lead to HLA alleles with different propensities for immune presentation, affecting T cell activation, or may perturb the binding of transcription factors that control HLA transcription (Supplemental Table S3). For example, the SATB1 pathway was predicted to have reduced expression in latent individuals relative to HAT patients (z-score=-2.093, p=0.008) (Supplemental Table S3). This protein mediates the development of T cells by reordering the chromatin of MHC (HLA) genes and affecting subsequent translation (Kumar et al. 2007). Polymorphisms in the HLA locus may therefore affect chromatin remodeling by SATB1 and subsequent HLA expression and activity. The HLA/MHC locus has been shown to be involved in several diseases with latent infections, most notably during HIV infection (Hentges et al. 1992), but also parasitic diseases related to *T. brucei*, including *Leishmania* (Quinnell et al. 2003) and *T. cruzi* (Cruz-Robles et al. 2004). Similarly, our work suggests that HLA-typing may have a role in understanding the latent infection phenotype associated with *T. b. gambiense* infections and provide a prognostic tool to predict disease severity. MHC alleles have not been shown to associate with susceptibility in mouse models for trypanosomiasis (Levine and Mansfield 1981), although an association has been shown in cattle (Karimuribo et al. 2011). HLA-G has also been previously been shown to be predictive of disease development in serological positive suspects (Ilboudo et al. 2014; Gineau et al. 2016). While making subsequent experimental analysis with animal models difficult, this further highlights the importance of field studies and hypothesis-generating research to understand novel characteristics.

Although our genetic analysis has suggested that T cell activation is involved in determining disease outcome, it was ambiguous if the reduced levels observed in latent infections relative to HAT patients was a cause or simply the result of low blood parasitemia. We therefore utilized experimental infections in a mouse model for trypanosomiasis using a specific inhibitor of co-stimulation dependent T cell activation. This confirmed that reduced T cell activation during trypanosome infection can lead to lower parasitemia and improved host condition. This is an important validation of the hypothesis that T cell activation plays a central role in determining disease severity, which emerged from the transcriptome and exome study.

Our data suggests that decreased T cell activation associates with a latent phenotype, both in the field and in an experimental model, it is unclear from our data how T cell activation leads to higher parasitemia and greater symptom severity. A possible contributor is a subpopulation of IFNγ-secreting CD8+ T cells that has been shown to be activated during trypanosome infection (Olsson et al. 1991). While IFNγ and a Th1 immune environment is associated with control of parasitaemia early in infection (Hertz and Mansfield 1999; Shi et al. 2003; Hertz et al. 1998; Shi et al. 2006), elevated or sustained IFNγ production can lead to more severe symptoms after the first peak of parasitemia (Namangala et al. 2001; Baetselier et al. 2001). The importance of a switch from a Th1 to Th2 immune environment has also been shown to be critical in mice infected with *T. b. gambiense* and animals that maintain high levels of IFNγ develop more severe symptoms (Inoue et al. 1999). Lack of expansion in the IFNγ-secreting CD8+ T cell population, either through genetic polymorphisms or experimentally via CTLA4-Ig, may lead to reduced IFNγ activity and an immune environment better able to control the parasite over the long-term. Interestingly, the most significantly associating SNPs from our study were found specifically in a class I *HLA* gene involved in CD8+ T cell presentation. Several other factors have been shown to modify the severity of symptoms during trypanosome infection, such as the cytokine IL-27 that ameliorates pathogenicity caused by activated CD4+ T cells (Liu et al. 2015). It is therefore likely that many T cell associating factors act in concert to generate an asymptomatic or latent phenotype and further studies are required. Nevertheless, we hypothesize that differential T cell activation contributes to the range of disease severity phenotypes observed in trypanosome infections, with latent infections associating with a relatively reduced T cell activation and clinical cases associating with elevated T cell activation.

Finally, we have demonstrated that trypanosome-induced T cell activation may be an appropriate target for intervention to reduce symptoms and improve outcome. While directly reducing T cell activation in the animal model reduced parasitemia and improved animal condition, eliciting such a profound reduction in T cell activity is not desirable in humans due to the variety of infections patients are likely to encounter in sub-Saharan Africa. Instead, targeting the parasite molecules that induce T cell activation may serve as an effective alternative. Prior work has shown that epitopes found on parasite variable surface glycoprotein (VSG) can elicit a T cell response (Mansfield 1994) and vaccination with VSG GPI-anchor in an animal model can lead to reduced pathogenicity (Magez et al. 2002). Additionally, *T. brucei* possesses a protein, termed trypanosome lymphocyte triggering factor (TLTF), that can directly induce activation in IFNγ-releasing CD8+ T cells (Vaidya et al. 1997). This protein has also been demonstrated to be present in *T. b. gambiense* (Bakhiet et al. 1996). Interestingly, TLTF is predicted to interact directly with CD8 and polymorphisms in CD8 or class 1 HLA alleles may therefore impact activity (Olsson et al. 1993). Antibodies raised against TLTF suppress the release of IFNγ by *in vitro* cultured T cells, although the therapeutic potential of these antibodies has not been assessed (Hamadien et al. 1999). By targeting the parasite factors that elicit T cell activation and the expansion of the IFNγ-secreting CD8+ T cell population, it may be possible to elicit a protective immune environment that mirrors the latent infection phenotype. Our field and animal model data suggests that this will allow patients to control infection, improving outcome.

In summary, our study suggests that genes involved in regulating T-cell activation are important in determining human *T. b. gambiense* infection outcomes. This builds on prior work demonstrating an association with APOL1 using the same patient samples. Our study is also an example of successfully applying transcriptome and genetic hypothesis-generating research to identify loci and pathways associating with the development of a latent asymptomatic phenotype in a human disease. Importantly, we have verified this predicted association using an animal model to clearly demonstrate the relevance of identified factors.

## Methods

### Study Design

Participants were identified during surveys organized by the Guinean National Control Programme (NCP) between 2007 and 2011 from three foci in Guinea (Dubreka, Boffa, and Forecariah)(Camara et al. 2005) according to the WHO and National Control Program policies (Ilboudo et al. 2011). For each participant, 500ul of plasma were collected and two ml of blood were taken on PAXgen Blood RNA tubes (PreAnalytix). All samples were frozen in the field at -20°C and were then stored at -80°C at CIRDES (Centre International de Recherche-Développement sur l'Elevage en zone Subhumide) until use. For each individual, the plasma was used to perform the immune trypanolysis test (TL) to detect surface antigens specific for *T. b. gambiense* (LiTat 1.3, LiTat 1.5) (Jamonneau et al 2010). Of the total cohort, an age-matched selection of 14 clinical cases (characterized as Card Agglutination Test for Trypanosomiasis (CATT) positive and trypanosomes were also detected using a mini Anion Exchange Centrifugation Technique (mAECT), followed by microscopy and /or examination of cervical lymph node aspirates by microscopy when adenopathies were present), 14 latent infections (CATT plasma titre 1/8 or higher, TL positive, no trypanosomes detected by mean of mAECT and / or examination of cervical lymph node aspirate during a two-year follow-up), and 7 uninfected controls (negative to the CATT, negative to the mAECT and negative to TL) were used in this study (mean age 37 years ±17). RNAs were extracted from blood collected on PAXgen Blood RNA tubes using the PAXgen^®^ Blood RNA kit (PreAnalytiX) according to manufacturer’s instruction. Two of the HAT patients were determined to be related after sequencing, adding power to the subsequent PLINK analysis.

### Ethics approval and consent to participate

All samples were collected as part of medical survey and epidemiological surveillance activities supervised by the Guinea National Control Program. All participants were informed of the objectives of the study in their own language and signed an informed consent form. This study is part of two larger projects. The first is aimed at improving diagnosis for which approval was obtained from the WHO (Research Ethics Review Committee), Institut de Recherche pour le Développement (Comité Consultatif de Déontologie et d'Ethique), and the University of Glasgow (MVLS Research Ethics Committee) ethical committees. The second study is TrypanoGEN, part of the H3Africa consortium, aimed at identifying the underlying genetic determinants of human susceptibility/resistance to trypanosome infections (Ilboudo et al. 2017), and ethical approval was sought from all subjects under the H3Africa consortium guidelines.

### Transcriptome Analysis

Illumina paired-end sequencing libraries were prepared from PBMC RNA using an Illumina TruSeq stranded mRNA kit. Fragment length ranged from 100-500 bp, with a median length of 150 bp. Libraries were sequenced by standard procedure on the Illumina HiSeq platform to yield paired-sequence reads 75 bp in length. Reads were aligned to the complete human cDNA library (GRCh38.79) using bowtie2-2.2.5 (Langmead and Salzberg 2012). Transcript counts were extracted using a custom PERL script and differential expression and significance calculated using DeSeq2v2.11(Love et al. 2014) for R. The mean alignment rate per sample for paired reads to the reference genome was 87.94 %.

### Pathway Analysis

Pathway analysis was performed using Ingenuity Pathway Analysis (IPA) Software 346717M (Qiagen) with a dataset of differentially expressed genes (adjusted p value < 0.1), corresponding to approximately 1,000 transcripts. The dataset was analyzed using the IPA core analysis tool to identify both direct and indirect relationships, using all data sources and tissue types but limited to experimentally observed relationships among human patient data.

### SNP Association Analysis

Illumina reads from the transcriptome analysis were aligned to the human genome (GRCh38.79) using Bowtie2-2.2.5. The average read depth for coding regions was 22.12. A sequence pileup was created using SAMtools 1.2 and single nucleotide polymorphisms (SNPs) called for the entire population using BCFtools 1.2. SNPs were further filtered to create a final SNP dataset containing binary SNPs of at least BCFtools quality 20. Genes and genomic features containing polymorphic loci were annotated using SnpEff4.1G. Associating SNPs were identified using PLINK1.7 with the standard association test, adaptive permutation and adjustment for false discovery to generate empirical p-values. fastSTRUCTURE was used to establish any underlying structure in the genomic data for values of K up to 10(Raj et al. 2014). Posthoc statistical power was calculated using the formula:

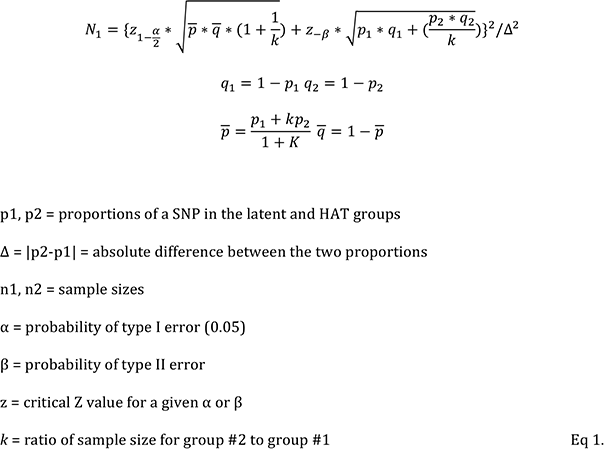

Sets of SNPs within gene-coding regions were grouped and additional set-based association tests were performed using a maximum of 10,000 permutations to find identify gene associations.

### Infection and strains

For the animal model of trypanosomiasis, 6-8 week old female BALB/c mice were housed in a specific pathogen-free environment and were used in accordance with HMG Home Office regulations. Bloodstream forms of the genome strain *Trypanosoma brucei* TREU927 were used for all assays (Gibson 2012). Mice were inoculated intraperitoneally with 10^4^ parasites. Parasitemia was assayed on each subsequent day using phase microscopy (Lumsden 1963).

### Flow cytometry

Single cells from blood and spleens were incubated with Fc block for 10 minutes before staining for 40 minutes with fluorochrome-conjugated antibodies raised against CD8a (APC anti-CD8a Biolegend #100712), CD4 (APC anti-CD4 Biolegend #100408), and several immune checkpoint proteins to assess activation (AlexaFluor488 anti-ICOS Biolegend #313514, PerCP5.5 anti-PD1 Biolegend #109120, and PE anti-LAG3 Biolegend #125208). Cells were washed, fixed with 4% paraformaldehyde, and analyzed on a MACSquant Flow Cytometer (Miltenyi). Subsequent population gating and analysis was performed using FlowJo software.

### *In vivo* treatments

CTLA4-Ig was provided by Bristol-Myers Squibb and administered by intraperitoneal injection at a dose of 10mg/kg every second day. The initial dose was administered at day -1 relative to inoculation. Human IgG1 was used as an isotype control in these studies.

## Data Access

The RNASeq datasets associated with the study are available in the EBI repository under the primary accession number PRJEB21742.

## Acknowledgements

The authors would like to thank Prof. Eileen Devaney for advice in writing the manuscript and to staff at Glasgow Polyomics for their support sequencing the samples. The sequencing and analysis undertaken in the project was funded by a Wellcome Trust Institutional Strategic Support Fund Catalyst Grant (097821/Z/11/B) and a Tenovus Scotland Project Grant (S14/18) awarded to PC and a Wellcome Trust Senior Fellowship awarded to AM (095201/Z/10/Z). Funding for the Wellcome Centre for Molecular Parasitology is provided by the Wellcome Trust (085349). The funders had no role in study design, data collection and interpretation, or the decision to submit the work for publication.

## Supplemental File Legends

**Supplemental Table S1. Transcripts that differ significantly between symptomatic and latent infections.** Transcripts significantly up in latent individuals (or significantly down in symptomatic cases), with an FDR-adjusted p value of <0.05 determined using DeSeq 2.0, are highlighted in blue. Transcripts significantly up in symptomatic cases (or significantly down in latent individuals) are highlighted in red.

**Supplemental Table S2. Disease or functional pathways that differ significantly differ between symptomatic and latent infections.** Pathways listed have p values of <0.05 determined using Ingenuity Pathway Analysis software. Inhibition (z-score < -2.0) or activation (z-score > 2.0) of a pathway in symptomatic HAT patients relative to latent infections is indicated. Molecules contributing to the pathway prediction are listed.

**Supplemental Table S3. Predicted upstream regulators of the pathway differences observed between latent individuals and symptomatic cases.** Regulators listed have p values of <0.05 determined using Ingenuity Pathway Analysis software. Inhibition (z-score < -2.0) or activation (z-score > 2.0) of each regulator in latent individuals relative to symptomatic cases is indicated.

**Supplemental Table S4. SNPs that significantly segregated between clinical and latent infections.** SNPs listed have an FDR-adjusted significance value of p < 0.05, assessed using PLINK. The alleles and proportions of each are shown for symptomatic cases and latent individuals. The odds ratio for the SNP to associate with the latent phenotype, and the resulting mutation type and gene are also shown.

**Supplemental Table S5. SNPs that significantly segregated between clinical and latent infections, grouped by gene.** SNPs were allocated to gene sets groups based on their position and analyzed using PLINK. This revealed genes that significantly segregated between clinical and latent infections with a nominal significance value of p < 0.05. The number of SNPS in each set, including non-significant and non-significant SNPS (including position) are listed.

**Supplemental Figure S1. ICOS, PD1, and LAG3 expression (markers of activation) on CD4+ T cells during infection.** The expression of ICOS, PD1, and LAG3 (markers of activation) on CD4+ T cells and ICOS and LAG3 on CD8+ T cells on days 5 and 11 of experimental TREU927 trypanosome infection in mice (n=5 per group, per time point), assessed via flow cytometry. This demonstrates that CTLA4-Ig (ABA) reduces T cell activation during infection relative to an isotype control (ISO).

